# The DNA damage response regulates the oocyte pool in mammals

**DOI:** 10.1101/816488

**Authors:** Ana Martínez-Marchal, Maria Teresa Guillot, Mònica Ferrer, Anna Guixé, Montserrat Garcia-Caldés, Ignasi Roig

**Affiliations:** Unitat de Citologia i Histologia, Departament de Biologia Cel·lular, Fisiologia i Immunologia, Facultat de Biociències, Universitat Autònoma de Barcelona, Cerdanyola del Vallès, Spain; Grup d’Inestabilitat i Integritat del genoma, Institut de Biotecnologia i Biomedicina, Universitat Autònoma de Barcelona, Cerdanyola del Vallès, Spain; Unitat de Biologia Cel·lular i Genètica Mèdica, Facultat de Medicina, Universitat Autònoma de Barcelona, Cerdanyola del Vallès, Spain

**Author notes:** Correspondence: I.R.

**Keywords:** Oocyte reserve, DNA damage response, CHK2, cyst breakdown, oogenesis, atresia

## Abstract

Mammalian oogonia proliferate without completing cytokinesis producing germ cell cysts. Within these cysts, oocytes differentiate and enter meiosis, promote genome-wide double-strand break (DSBs) formation which repair by homologous recombination leads to synapsis of the homologous chromosomes. Errors in homologous recombination or synapsis trigger the activation of surveillance mechanisms, traditionally called ‘pachytene checkpoint’, to either repair them or send the cells to programmed death. Contrary to what is found in spermatocytes, most oocytes present a remarkable persistence of unrepaired DSBs at pachynema. Simultaneously, there is a massive oocyte death accompanying the oocyte cyst breakdown. This oocyte elimination is thought to be required to properly form the follicles, which constitute the pool of germ cells females will use during their adult life. Based on all the above mentioned, we hypothesized that the apparently inefficient meiotic recombination occurring in mouse oocytes may be required to eliminate most of the oocytes in order to regulate the oocyte number, promote cyst breakdown and follicle formation in mammalian females. To test this idea, we analyzed perinatal ovaries to evaluate the oocyte population, cyst breakdown and follicle formation in control and mutant mice for the effector kinase of the DNA damage response, CHK2. Our results confirm the involvement of CHK2 in the elimination of oocytes that accumulate unrepaired DSBs and show that CHK2 regulates the number of oocytes in fetal ovaries. We also show that CHK2 is required to eliminate oocytes as a result of LINE-1 activation, which was previously shown to be responsible for fetal oocyte loss. Nonetheless, the number of oocytes found in *Chk2* mutant ovaries three days after birth was similar to that of control ovaries, suggesting the existence of CHK2-independent mechanisms capable of eliminating oocytes. *In vitro* inhibition of CHK1 rescued the oocyte number in *Chk2* mutant ovaries suggesting that CHK1 regulates postnatal oocyte death. Moreover, both CHK1 and CHK2 functions are required to timely breakdown cyst and form follicles. Altogether, we propose the DNA damage response controls the number of oocytes present perinatally and is required to properly break down oocyte cysts and form follicles, highlighting the importance of the DNA damage response in setting the reserve of oocytes each female will use during their entire lifespan.

## Introduction

Meiosis is the reductional division of the genome that generates haploid cells. At the onset of meiotic prophase I, SPO11 creates double-stranded breaks (DSBs) all over the genome, which repair by homologous recombination leads to the pairing and synapsis of the homologous chromosomes (Baudat et al., 2000). These processes are tightly regulated to avoid possible deleterious effects originated by errors in recombination or synapsis. When errors are produced, meiocytes delay their cell cycle progression, and can even activate programmed cell death (Subramanian and Hochwagen, 2014). Two mechanisms have been identified to lead mammalian meiocytes into apoptosis. One dependent on DSB formation and recombination progression, and another independent of SPO11-generated DSBs (Barchi et al., 2005; Bolcun-Filas et al., 2014; Di Giacomo et al., 2005; Marcet-Ortega et al., 2017; Pacheco et al., 2015). Studies in spermatocytes and oocytes have revealed the central role of the DNA damage response (DDR) effector kinase, CHK2, has in the arrest of mammalian meiocytes with persistent recombination intermediates (Bolcun-Filas et al., 2014; Pacheco et al., 2015).

Mammalian gametogenesis is characterized by a marked sexual dimorphism (Morelli and Cohen, 2005). One of the things that have intrigued researchers in the field is the distinct control of meiotic recombination that occurs in oocytes and spermatocytes (Lenzi et al., 2005; Roig et al., 2004). We and others reported that while most human spermatocytes have repaired most DSBs at pachynema, the vast majority of human oocytes still present multiple unrepaired DSBs at this stage.

Another distinct characteristic of oogenesis is that most of the oocytes formed will eventually die (Hunter, 2017). In humans, it is believed that only ∼10% of the oocytes that initiate meiosis will end up forming a follicle (Baker, 1963). In mice, there is a massive oocyte elimination around birth when follicles form. Oogonium, before differentiating into oocytes, go through several mitotic divisions with incomplete cytokinesis, thus, resulting in a syncitium of cells, called cyst. Therefore, when oocytes enter meiosis, they are connected to their sister cells. At the end of the meiotic prophase, cysts break down so single oocytes can be surrounded by stromal ovarian cells to form follicles, which will be the pool of germ cells females will use in their reproductive lifetime.

It is still unknown the reason behind the effort of eliminating the vast majority of the oocytes produced. it has been speculated it may help in disassembling the oocyte cysts to separate individual oocytes, so follicles can form (Pepling, 2012). Also, oocytes in cyst are able to transfer organelles to neighboring connected oocytes, in a process reminiscent of the Drosophila nurse cell dumping (Lei and Spradling, 2016). However, the importance of this process is not clear, since mutant mice defective for TEX14, which is the ring canal protein required to transfer the cytoplasmic content between cyst cells, are fertile (Greenbaum et al., 2009). Similarly, it is not known the mechanisms that regulate oocyte death in wild-type mice. Apart from recombination and synaptic errors, the activation of LINE-1 during fetal development can also trigger oocyte death (Malki et al., 2014). however, the mechanism behind it is also unknown (Hunter, 2017).

Based on the high levels of unrepaired DSBs present in wild-type oocytes at pachynema, we hypothesized that part of the fetal oocyte death is due to the activation of the CHK2-dependent DDR in the oocytes that accumulate more unrepaired DSBs. We also hypothesized that the DDR-dependent oocyte elimination would contribute to proper cyst breakdown and follicle formation. To test these ideas, we analyzed the number of oocytes, cysts breakdown and follicle formation in perinatal female ovaries from control and *Chk2*^-/-^ mice. Our results confirm the involvement of CHK2 in the elimination of oocytes that accumulate unrepaired DSBs and show that CHK2 regulates the oocyte number in fetal ovaries. We also show that CHK2 is required to eliminate oocytes as a result of LINE-1 activation, suggesting that the mechanism behind LINE-1-triggered oocyte death depends on the DDR. Nonetheless, control and *Chk2* mutant samples had the same number of oocytes at 4 days postpartum (dpp). Our results also provide evidence that the CHK1 function is required to eliminate *Chk2*^*-/-*^ oocytes after birth. Moreover, cyst breakdown was also disrupted when the two effector kinases of the DDR where inhibited, linking the regulation of the oocyte population with cyst breakdown. Altogether, our data show that the DDR regulates the oocyte pool in mammals and allows proper cyst breakdown and follicle formation.

## Results

### *Chk2* mutant fetal ovaries accumulate oocytes with more unrepaired DSBs than controls

Our model predicts that in the absence of CHK2 ovaries will accumulate oocytes with high numbers of unrepaired DSBs that would normally be eliminated. To test this, we counted the number of γH2AX patches, as a surrogate of unrepaired DSBs (Pacheco et al., 2019; Roig et al., 2004), in control and *Chk2* mutant oocytes at different stages of meiotic prophase. As mentioned earlier and described elsewhere (Pacheco et al., 2019; Roig et al., 2004), wild-type oocytes at pachynema presented multiple unrepaired DSBs. Importantly, *Chk2* mutant pachytene-stage oocytes had significantly more patches of γH2AX than control cells (p<0.0001 t-tests, Fig. 1 and Table S1). This finding corroborates our model and suggests that CHK2 eliminates the oocytes with a higher number of unrepaired DSBs at pachynema. Nonetheless, as oocytes completed meiotic prophase the difference of unrepaired DSBs between the two genotypes decreased. At late diplonema, mutant and control oocytes displayed similar numbers of γH2AX patches (p>0.05 t-test, Fig. 1 and Table S1). This result could be a manifestation of the ability of the oocytes to repair the damage past the pachytene stage, as it has been proposed before (Bolcun-Filas et al., 2014). Alternatively, it could also be interpreted as oocytes accumulating unrepaired DNA damage could be eliminated by CHK2-independent mechanism occurring after the pachytene stage.

**Figure 1.**
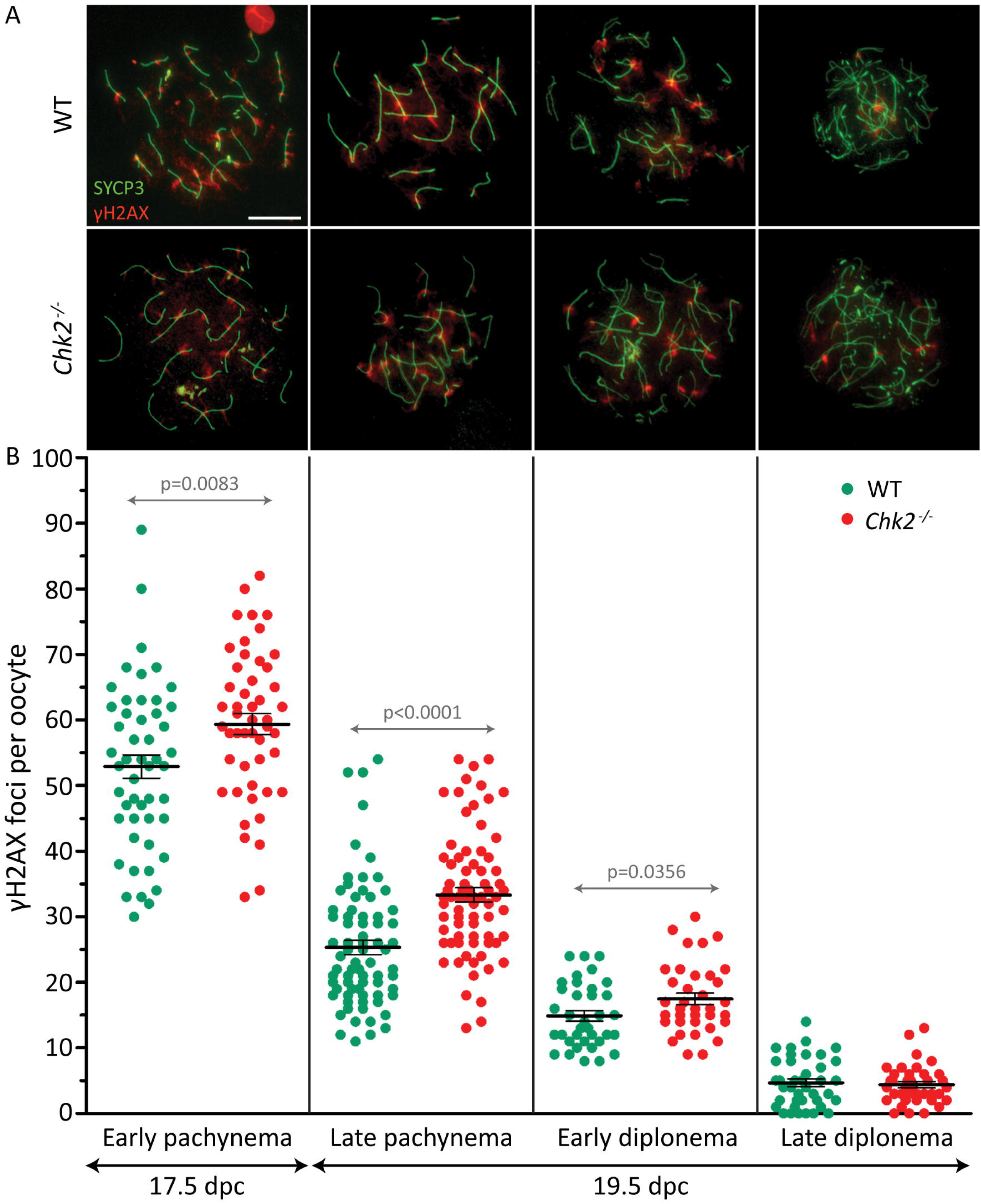
*Chk2*^*-/-*^ oocytes present a higher number of unrepaired DSBs at pachynema and early diplonema than controls. (A) Representative images of control (WT) and *Chk2*^*-/-*^ oocytes at pachynema and diplonema. The cells are immunostained against SYCP3 (green) and ?H2AX (red). Scale bar represents 10 μm and applies to all images. (B) Quantification of the number of γH2AX patches found in WT and *Chk2*^*-/-*^ oocytes at pachynema and diplonema. The horizontal lines represent the means ± standard error of the mean (SEM).

### Perinatal *Chk2* mutant ovaries have more oocytes than control ovaries

To test our model, we counted the number of oocytes present in control and *Chk2* mutant ovaries from 15.5 dpc to 4 dpp (22.5 dpc) mice. We expected that *Chk2* mutant ovaries would present more oocytes than wild-type samples after most oocytes have reached the pachytene stage, around 17.5-19.5 dpc. To perform this analysis, we serially sectioned the ovaries and immunostained every other section against the germ cell marker DDX4 (Fig. 2A, B). Our control mice showed the expected pattern: at 15.5 dpc, ovaries contained the maximum number of germ cells, on average almost 9000 oocytes (Fig. 2C and Table S2). At 17.5 dpc, half of the control oocytes had been eliminated and the ovaries displayed an average number of 4207 oocytes. From this moment, the number of oocytes found in our control mice tended to slowly decrease until 4 dpp. As expected, *Chk2* mutant ovaries displayed a similar number of oocytes than control samples at 15.5 dpc (Fig. 2C and Table S2). However, the number of oocytes found at 17.5 dpc in *Chk2* mutant samples was significantly higher than the one found in control samples (p=0.0001, T test). These data suggest that CHK2 is required to eliminate the fetal oocytes around 16.5 dpc, as our model predicted. Unexpectedly, the number of oocytes present in *Chk2* mutant ovaries declined significantly during the following days, until it matched the number of oocytes found in wild-type samples. These results suggest that there must be a CHK2-independent mechanism that is able to eliminate *Chk2*^*-/-*^ oocytes after birth.

**Figure 2.**
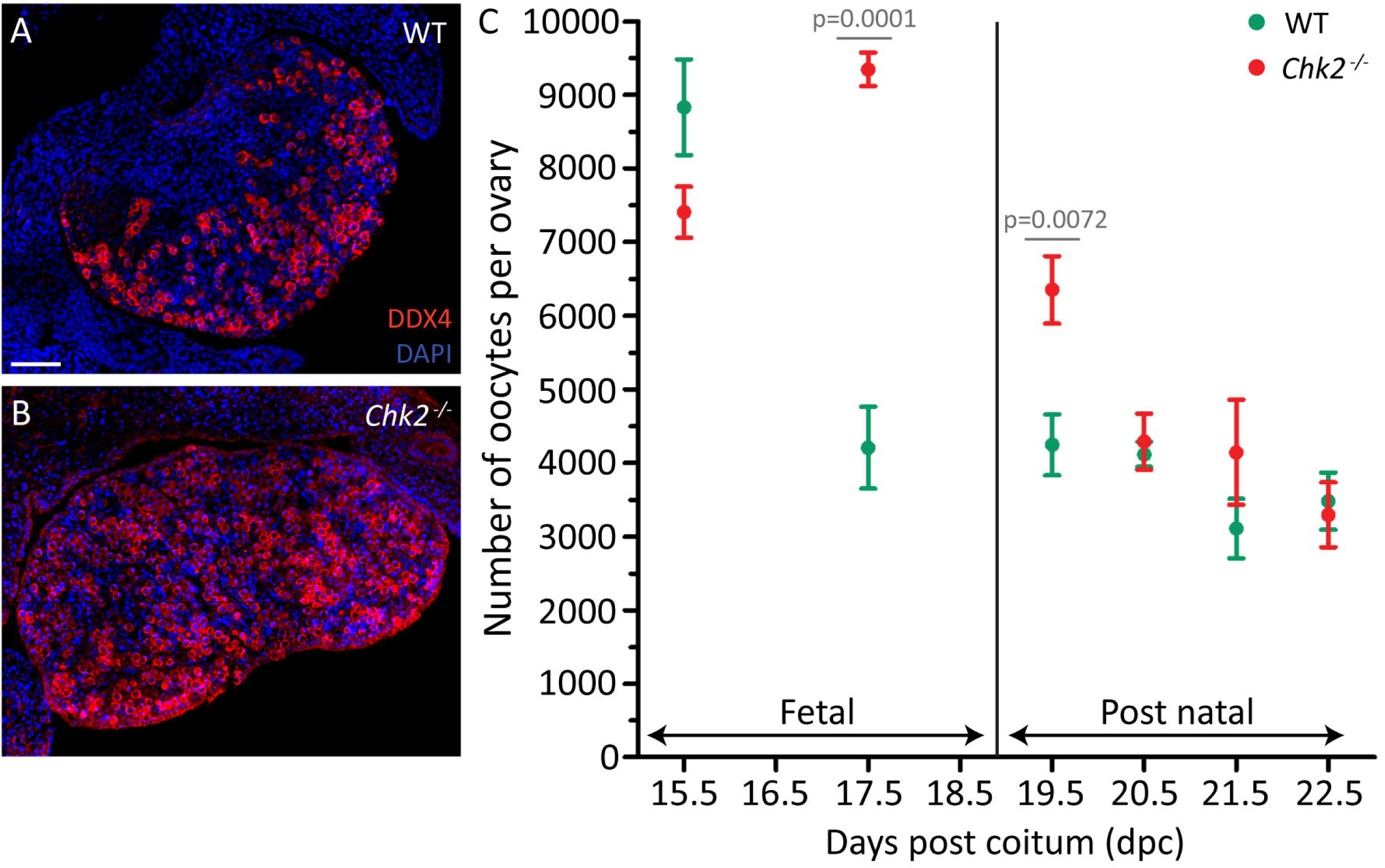
CHK2 regulates the number of oocytes in fetal ovaries. (A-B) Representative images of 17.5 dpc control (WT, A) and *Chk2*^*-/-*^ (B) histological sections of ovaries immunostained against DDX4 (red) and the DNA counterstained with DAPI (blue). Scale bar represents 40 μm and applies to both images. (C) Number of oocytes in fetal (15.5-17.5 dpc) and postnatal (19.5-22.5 dpc) control (WT) and *Chk2*^*-/-*^ ovaries. The round symbols represent the mean and the lines the SEM.

### Lack of CHK2 does not compromise cyst breakdown and follicle formation

Our model also predicts that *Chk2* mutant ovaries will present defects in cyst breakdown and follicle formation. Thus, we classified the oocytes from control and mutant ovaries from 15.5 dpc to 4 dpp into three categories: oocytes in cyst (those sharing cytoplasm with their neighbors, Fig. 3A), single oocytes (in which the limit of the oocyte cytoplasm was clearly visible, Fig. 3B) and oocytes in follicles (in which the oocytes were surrounded by follicular cells; Fig. 3C). At 15.5 dpc, approximately 40% of the oocytes from control and mutant ovaries remained in cysts, while the rest were already individualized (Fig. 3D and Table S3). At 17.5 dpc, fifty percent of the control oocyte population had disappeared. Remarkably, this translated into the loss of 50% of the oocytes that were forming cysts, but also of those that were already individualized. As expected, 15.5 dpc and 17.5 dpc *Chk2* mutant ovaries contained similar number of oocytes in cysts and individual oocytes, similarly to what was found in control and mutant samples at 15.5 dpc. These data suggest that the massive CHK2-dependent oocyte elimination occurs in both oocytes in cysts and individual oocytes. Follicles began to appear in control samples at 17.5 dpc and formed most of the oocyte pool from 3 dpp (21.5 dpc) onwards. Follicle formation in *Chk2* mutants was similar to the one found in controls, follicles first appeared in newborn mice and constituted most of the oocyte pool from 3 dpp onwards. These data suggest that CHK2 is dispensable for proper cyst breakdown and follicle formation.

**Figure 3.**
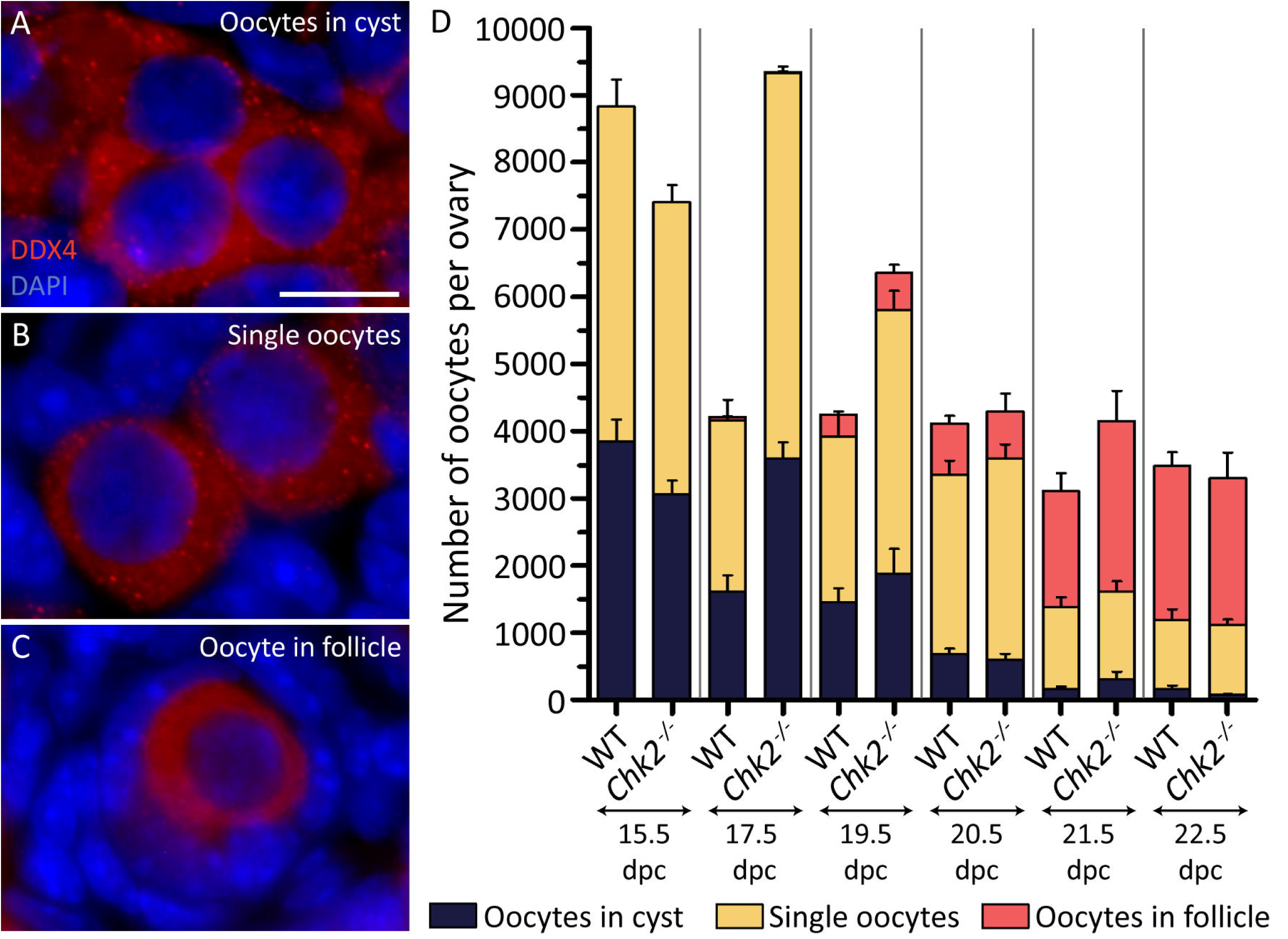
CHK2 seems to be dispensable for cyst breakdown and follicle formation. (A-C) Representative images of oocytes immunostained against the germ cell marker DDX4 forming a cyst (A), two single oocytes (B) and forming a follicle (C). The DNA is counterstained with DAPI (blue). Scale bar represents 10 μm and applies to all images. (D) Number of oocytes classified in the three different types described above in 15.5-22.5 dpc ovaries. The lines represent the mean ± SEM.

### *Chk2* mutation partially rescues the oocyte loss occurring in *Spo11* mutants

At the onset of meiotic prophase, SPO11 generates multiple DSBs that will drive meiotic recombination and homologous chromosomes synapsis (Baudat et al., 2000). Thus, based on our model, we expected that the massive perinatal oocyte loss that occurs in *Spo11* mutants would be independent of CHK2. To test this, we counted the number of oocytes present in newborn ovaries from *Spo11*^*-/-*^ and *Spo11*^*-/-*^ *Chk2*^*-/-*^ mice. *Spo11*^*-/-*^ ovaries presented approximately 25% of the number of oocytes found in *Chk2*^*-/-*^ mice of the same age (p<0.0001, T test, Fig. 4 and Table S4). Significantly, *Spo11*^*-/-*^ *Chk2*^*-/-*^ ovaries contained twice as many oocytes as *Spo11*^*-/-*^ ovaries, suggesting that CHK2 was responsible for part of the oocyte death occurring in *Spo11*^*-/-*^ ovaries (p=0.0118; T test). *Spo11*^*-/-*^ oocytes have been reported to present DNA damage at pachynema (Carofiglio et al., 2013; Rinaldi et al., 2017). Thus, CHK2 may be activated as a response to the presence of this DNA damage and ultimately this leads to the elimination of some of these oocytes, as previously reported (Rinaldi et al., 2017). Importantly, *Spo11*^*-/-*^ *Chk2*^*-/-*^ ovaries contained half the number of oocytes found in *Chk2*^*-/-*^ ovaries (p=0.0022; T test). These results suggest that CHK2-independent mechanisms are also responsible for part of the oocyte death occurring in *Spo11*^*-/-*^ fetal ovaries.

**Figure 4.**
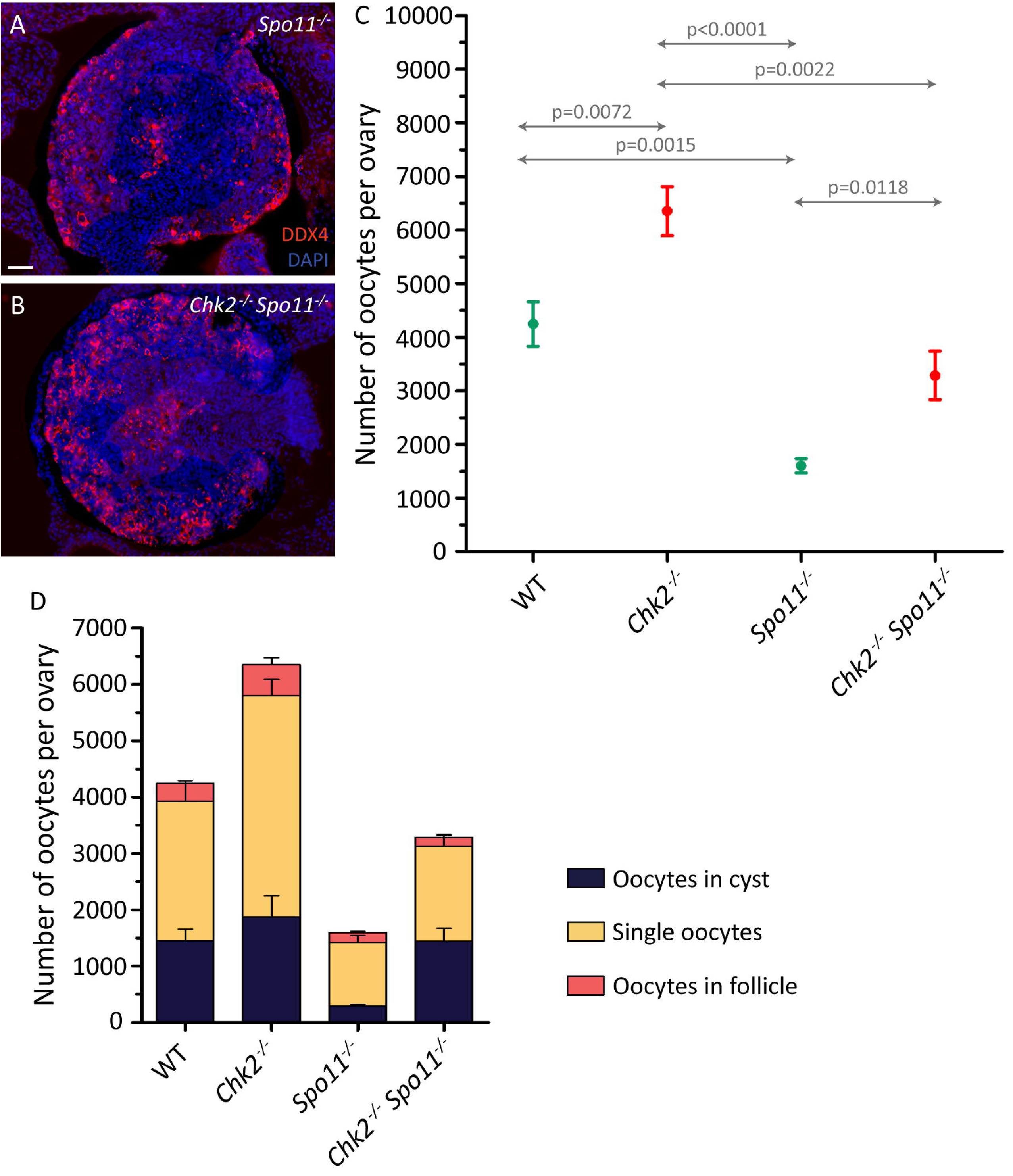
CHK2 rescues the number of oocytes in cyst in a SPO11 background. (A-B) Representative images of 19.5 dpc *Spo11*^*-/-*^ (A) and *Chk2*^*-/-*^ *Spo11*^*-/-*^ (B) histological sections of ovaries immunostained against DDX4 and counterstained with DAPI. Scale bar represents 40 μm and applies to both images. (C) Number of oocytes in 19.5 dpc control (WT), *Chk2*^*-/-*^, *Spo11*^*-/-*^ and *Chk2*^*-/-*^ *Spo11*^*-/-*^ ovaries. The round symbols represent the mean and the lines the SEM. The data for control and *Chk2*^*-/-*^ samples were taken from Figure 2. (D) Number of oocytes classified in the three different types (oocytes in cyst, single oocytes, and oocytes in follicle) in 19.5 dpc control (WT), *Chk2*^*-/-*^, *Spo11*^*-/-*^ and *Chk2*^*-/-*^ *Spo11*^*-/-*^ ovaries. The lines represent the mean ± SEM. The data for control and *Chk2*^*-/-*^ samples were taken from Figure 3.

Moreover, removal of CHK2 in *Spo11* mutant oocytes selectively rescued oocytes in cysts (Fig. 4D and Table S4), suggesting that CHK2 participates in the cyst breakdown when SPO11 function is suppressed.

### CHK1 function is required to eliminate oocytes in *Chk2*^*-/-*^ ovaries in vitro

Contrary to what our model predicted, the number of oocytes present in *Chk2*^*-/-*^ ovaries significantly declined after birth, suggesting the existence of an alternative mechanism eliminating oocytes that accumulate DNA damage. The DDR relies on the activation of two effector kinases, CHK1 and CHK2, in order to repair the DNA damage, arrest cell cycle progression, and, if necessary, induce apoptosis (Stracker et al., 2009). Thus, we wondered if CHK1 was compensating for the loss of CHK2 and hence, was responsible for the postnatal elimination of *Chk2*^*-/-*^ oocytes. To test this hypothesis, we established an organotypic culture that allowed follicle formation *in vitro* (Fig S1), and cultured newborn *Chk2*^*-/-*^ ovarian samples in presence of different concentrations of LYS2603618, a specific CHK1 inhibitor (CHK1i) (Dai and Grant, 2010). We observed no difference in the number of oocytes found in cultured ovaries under a low concentration of CHK1i (1 μM) compared to the DMSO controls (P>0.05, T test; Fig. 5, Table S5). However, the addition to the culture medium of 5μM LYS2603618 resulted in an increased presence of oocytes after culture (p=0.0473, T test; Fig. 5). Significantly, these samples contained three times more oocytes in cyst and twice as many individual oocytes than the DMSO-treated ones (p=0.0049 and p=0.0081 respectively; T test; Fig. 5, Table S5). Also, follicle formation seemed to be reduced in these treated samples, although this difference was not statistically significant (p=0.2687, T test). These data suggest that CHK1 eliminates *Chk2*^*-/-*^ oocytes after birth and helps to breakdown the oocyte cysts to form follicles.

**Figure 5.**
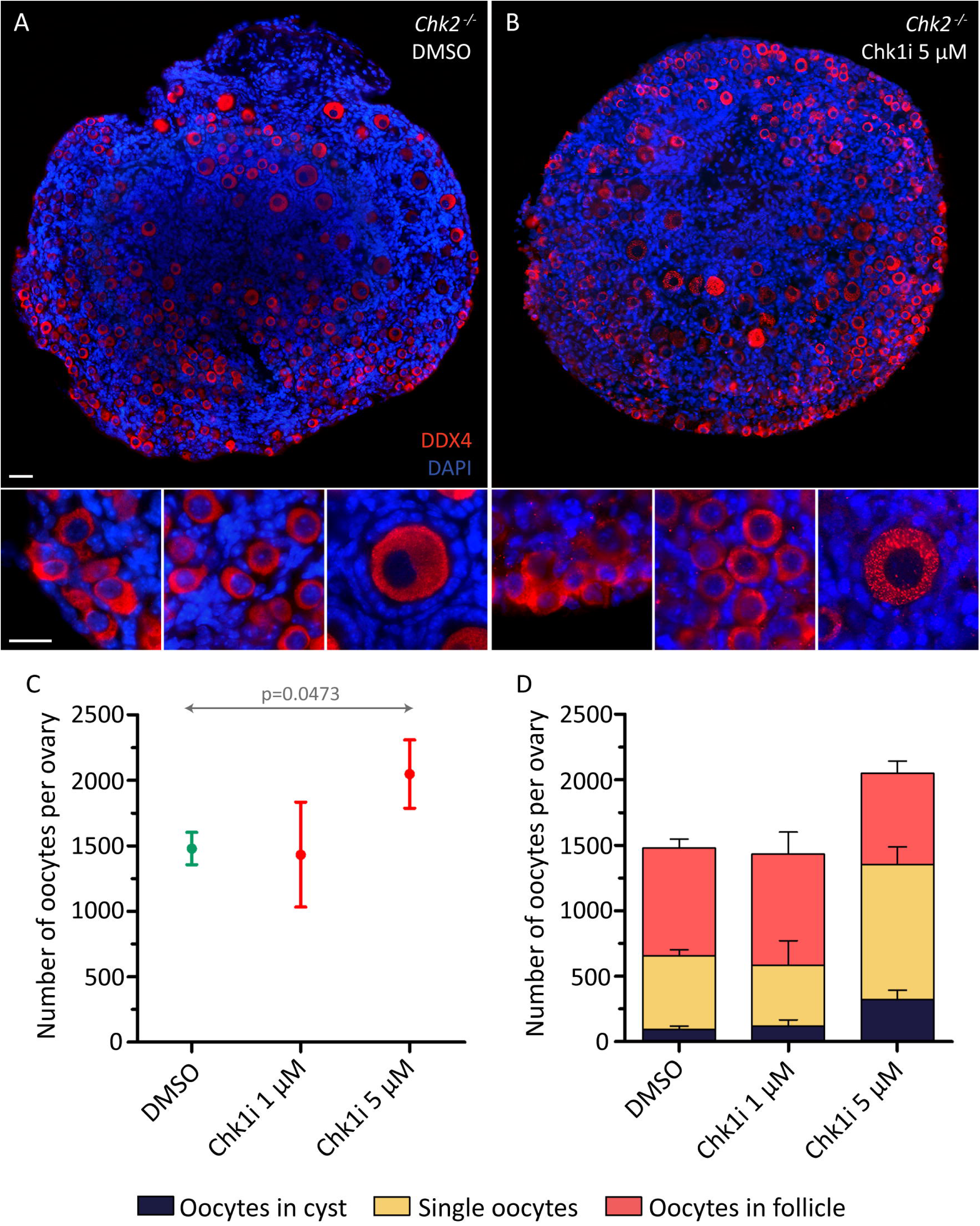
The inhibition of CHK1 *in vitro* rescues the oocyte number in *Chk2*^*-/-*^ ovaries. (A-B) Histological sections of *Chk2*^-/-^ DMSO-treated ovaries (A) and 5 μM LY2603618 (CHK1i)-treated ovaries (B) immunostained against DDX4 and counterstained with DAPI. The inserts show the detail of oocytes in cyst (left), single oocytes (center) and a follicle (right). The scale bar in the top image represents 40 μm and applies to both top images. The scale bar in the insert represents 20 μm and applies to all bottom images. (C) Number of oocytes found in *Chk2*^*-/-*^ 19.5 dpc ovary after 5 days of culture exposed to DMSO or 1 µM or 5 µM of CHK1 inhibitor. The round symbols represent the mean and the lines the SEM. (D) Number of oocytes classified in the three different types (oocytes in cyst, single oocytes and oocytes in follicle) for the same ovaries. The lines represent the mean ± SEM per each category.

### The oocyte death mediated by LINE-1 activation depends on CHK2

The activation of the transposable element LINE-1 triggers fetal oocyte death (Malki et al., 2014). Since LINE-1 retrotransposition into the genome may cause DNA damage (Belgnaoui et al., 2006), we wondered if the fetal oocyte death caused by LINE-1 would depend on the activation of the DDR. To test this, we treated 11.5 dpc pregnant mice carrying either wild type or *Chk2*^*-/-*^ fetuses for six days with Azidothymidine (AZT) and collected fetal ovaries at 17.5 dpc. AZT is a nucleoside analog that specifically inhibits the retrotranscriptase activity required for LINE-1 retrotransposition into the genome. When we tested the previously reported concentration (5mg/Kg, Malki et al., 2014) on wild type mice, we were unable to observe an increase in the number of oocytes present in the AZT-treated ovaries (Fig. 6, Table S6). However, when we raised the dose to 15 mg/kg, the AZT-treated ovaries contained significantly more oocytes than water controls (Fig. 6E, Table S6, p=0.0090, T test). Interestingly, the AZT-treated wild-type ovaries had less oocytes than *Chk2*^*-/-*^ controls, treated with water (p=0.0421, T test), suggesting that *Chk2* mutation could rescue more oocytes than AZT inhibition. When we analyzed the effect of the AZT treatment on the *Chk2* mutants we did not observe any difference in the number of oocytes recovered in the control or in the AZT-treated samples, at any of the doses used (P>0.05, T test). These data show that the oocyte decline caused by the activation of LINE-1 during fetal development is dependent on CHK2. Similarly to what we observed in the *Chk2*^*-/-*^ ovaries, treatment with AZT had no influence in the cyst breakdown and follicle formation in neither wild type or *Chk2* mutant samples (Fig 6F and Table S6). These data reinforce the similarities between the effect of the *Chk2* mutation and the treatment with AZT and further supports the idea that the oocyte death caused by LINE-1 activation is dependent on the DDR function.

**Figure 6.**
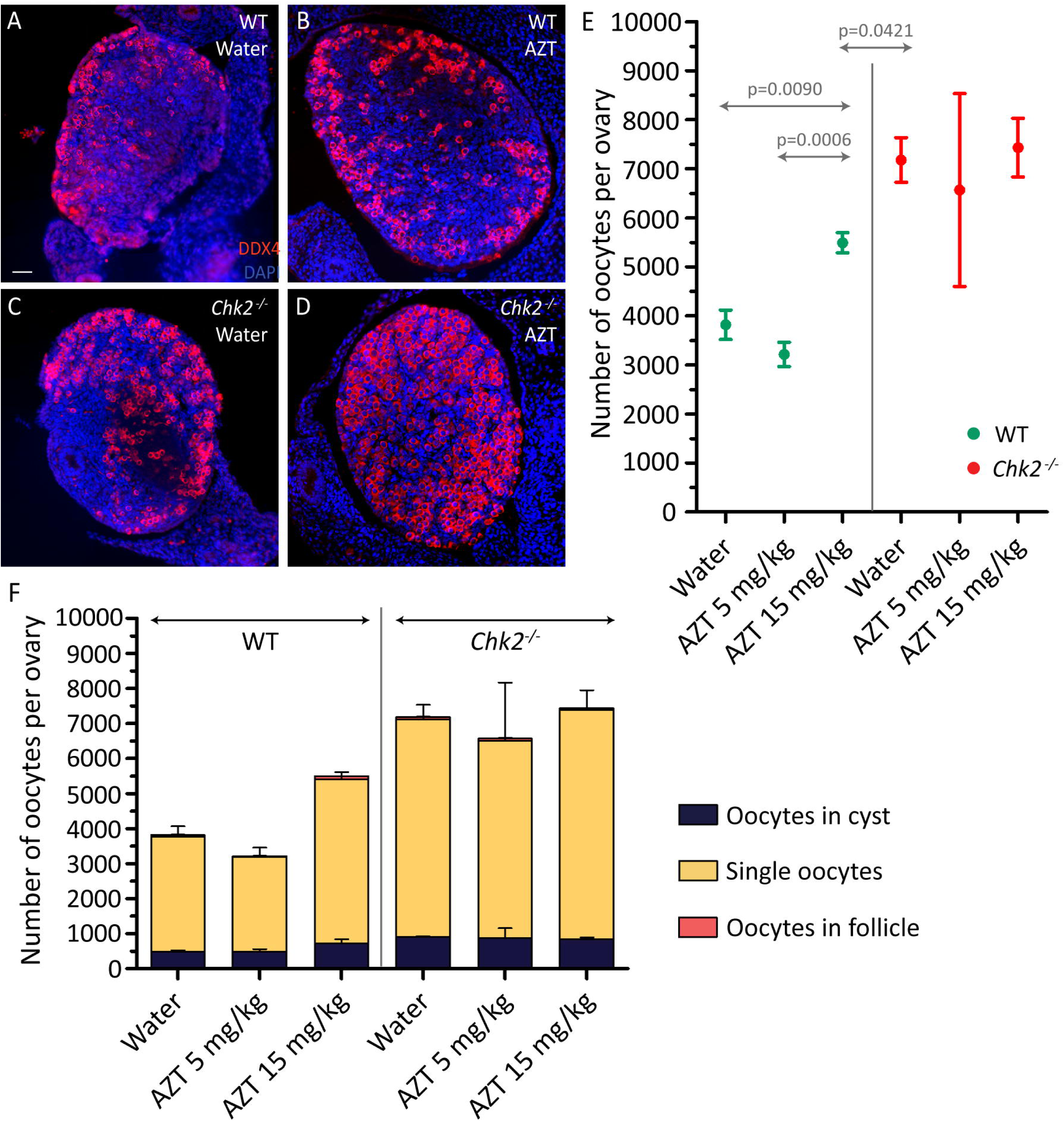
LINE-1 inhibition rescues the oocyte number in WT ovaries, but not in *Chk2*^*-/-*^ovaries. (A-D) Histological sections of Control (WT) treated with water (A), AZT-inhibited control (B), *Chk2*^-/-^ treated with water (C) and AZT-inhibited *Chk2*^*-/-*^ (D) ovaries immunostained against DDX4 and counterstained with DAPI. Scale bar represents 40 μm and applies to all images. (E) Number of oocytes per ovary after 5 days of water or AZT treatment. The round symbols represent the mean and the lines the SEM. (F) Number of oocytes classified in the three different types (oocytes in cyst, single oocytes and oocytes in follicle) for the same ovaries. The lines represent the mean ± SEM per each category.

## Discussion

The extent of the mammalian female reproductive lifespan is mostly defined by the number of oocytes that remain in the ovaries after birth. In this study, we have uncovered the critical role of the DDR in controlling the oocyte pool in mammals. Our data show that oocytes that accumulate unrepaired DSBs at pachynema are eliminated by the CHK2-dependent DDR during fetal development. Moreover, the existence of a CHK1-dependent mechanism eliminating *Chk2* mutant oocytes evidences the importance of the DDR in regulating mammalian oogenesis.

Mammalian gametogenesis is sexually dimorphic in several aspects (Morelli and Cohen, 2005). One of them is the apparently different efficiency in completing meiotic recombination of spermatocytes and oocytes (Lenzi et al., 2005; Roig et al., 2004). While mammalian spermatocytes complete repair of most of their DSBs at pachynema; oocytes show multiple unrepaired DSBs at this stage (Roig et al., 2004). Our findings suggest that the existence of these unrepaired DSBs is crucial to set the oocyte population during fetal development. Thus, the DDR is at least partly responsible to determine the perinatal oocyte pool.

Noteworthy, our data show that this CHK2-mediated meiotic response to DNA damage can react to SPO11-originated DSBs as well as other sources of DNA damage. It has been previously reported that *Spo11*^*-/-*^ oocytes present markers of DNA damage (Carofiglio et al., 2013; Rinaldi et al., 2017), and these can activate a CHK2-dependent oocyte elimination (Rinaldi et al., 2017). Our results confirm these findings revealing that the DDR is partly responsible for eliminating *Spo11* mutant oocytes. However, the observed rescue in 1 dpp *Spo11*^*-/-*^ *Chk2*^*-/-*^ ovaries does not reach *Chk2*^*-/-*^ numbers, as one would expect if the DDR activation was the only mechanism eliminating *Spo11*^*-/-*^ oocytes. In fact, our data show that the DDR is not the main mechanism to eliminate *Spo11*^*-/-*^ oocytes during fetal development since only ∼35% of the oocytes are eliminated by a CHK2-dependent pathway. Thus, other mechanisms should be accounted for the elimination of most of the *Spo11* mutant fetal oocyte pool. Presumably, the response to the presence of unsynapsed chromosomes may be responsible for this cell death. At the end of meiotic prophase, unsynapsed chromosomes are silenced by the activation of a cascade of events involving several key proteins of the DDR, such as ATR, MDC1 or H2AX that ultimately leads to their silencing (Ichijima et al., 2011, 2012; Mahadevaiah et al., 2008; Turner et al., 2005). This mechanism, known as meiotic silencing of unsynapsed chromosomes (MSUC), has been shown to be sufficient to eliminate oocytes with asynaptic chromosomes (Cloutier et al., 2015). Thus, it is plausible that most of the *Spo11*^*-/-*^ fetal oocytes are eliminated as a consequence of the MSUC mechanism. This finding highlights the existence of different surveillance mechanisms that monitor meiotic prophase progression which are activated by different events, such as asynapsis or presence of recombination intermediates (Di Giacomo et al., 2005). Nonetheless, the fact that part of the signaling machinery that participates in the response to synapsis and recombination defects is shared by both pathways (such as, ATR (Pacheco et al., 2018; Royo et al., 2013; Widger et al., 2018)) makes the study of these mechanisms very complex.

Apart from synaptic and recombination defects, the activation of LINE-1 retrotransposon can also trigger fetal oocyte death (Hunter, 2017). During germ cell development, LINE-1 is derepressed as a consequence of the erasing of the genome methylation marks to initiate the epigenetic reprogramming of the future egg. Restoration of DNA methylation will not take place after birth. Thus, meiotic prophase oocytes suffer from an increase in the expression of LINE-1 (Malki et al., 2014). This raise in LINE-1 activation is associated with the loss of approximately 50% of the oocytes, occurring between embryonic days 15.5 and 17.5. To date, it is not clear what mechanism is associating LINE-1 activation to oocyte death (Hunter, 2017). Since LINE-1 incorporation into the genome can cause DNA damage and the inhibition of LINE-1 activity was reported to cause an oocyte rescue similar to the one we observed in *Chk2* mutants (Hunter, 2017; Malki et al., 2014), we investigated if CHK2 was required for the oocyte loss caused by LINE-1 activity. We found problems recapitulating the previously published results (Malki et al., 2014), and had to triplicate the dose of AZT administered in our wild type mice to observe a rescue of oocytes. Importantly, this rescue was not complete, as it was previously reported. The observed differences may be attributable to the different genetic backgrounds that were used in the two studies. Nonetheless, it evidenced that LINE-1 activation was responsible for at least part of the fetal oocyte death that occurs in wild-type mice. However, AZT treatment had no effect on *Chk2* mutant mice, suggesting that the oocyte death originated by LINE-1 was also dependent on CHK2. Based on these data we propose that the mechanism behind LINE-1-induced oocyte death is the formation of DNA damage as a result of LINE-1 transposable activity, which may ultimately lead to the activation of CHK2 and the elimination of the fetal oocytes.

Our data also reveal the involvement of CHK1 in the elimination of oocytes with persistent DNA damage. The study of the function of CHK1 during meiotic prophase has been challenging because CHK1 is required to allow embryo development in mammals (Liu et al., 2000). Furthermore, the use of conditional mutants has shown that CHK1 is also required for germ cell proliferation, at least in the male (Abe et al., 2018). Our *in vitro* approach allowed us to show that CHK1 is able to compensate for the loss of CHK2. This finding explains why the number of oocytes found in 4 dpp *Chk2*^*-/-*^ ovaries is the same as those in wild type ovaries. Unexpectedly, CHK1 activation seems to be delayed as compared to CHK2. This opens the possibility that during meiotic prophase CHK2 and CHK1 may be differentially regulated. Sensitivity to DNA damage differs significantly from early prophase to dictyate-arrested oocytes. While leptotene oocytes endure hundreds of DSBs, dictyate oocytes are very sensitive to DNA damage (Lenzi et al., 2005; Suh et al., 2006). This is partly accomplished by regulating the activity of TAp63α, a key protein of the DDR in oocytes (Kim and Suh, 2014). P63 is required to eliminate dictyate-arrested oocytes weakly irradiated (0.3 Gy; Suh et al., 2006). However, p63 is not expressed until oocytes reach mid meiotic prophase (17 dpc onwards) and its activity is actively inhibited (Kim and Suh, 2014), suggesting that p63 activity might be deleterious for meiotic prophase oocytes. Thus, it is plausible that different pathways of the DDR are selectively regulated during oogenesis in order to first allow meiotic DSB formation and repair to promote homologs synapsis and crossover formation. Once this is accomplished, the DDR may activate alternative, or supplementary, pathways to achieve a higher sensitivity to DNA lesions, in order to assure only high-quality oocytes can pass their genetic information to the next generation. Thus, we propose CHK2 may participate in the surveillance mechanisms that control the progression of DSB repair until mid pachynema. This mechanism is responsible for the elimination of approximately 50% of the oocytes during fetal oogenesis. Once this meiotic surveillance mechanism is met, an alternative or supplementary CHK1-dependent pathway may become active to ensure the genetic integrity of the remaining oocytes. This model would explain why CHK1 cannot compensate for the loss of CHK2 during fetal development but can do it later on, once the oocytes may have reached a particular stage (late pachynema, diplonema and/or dictyate).

The perinatal massive oocyte death that naturally occurs in mouse oogenesis has been historically associated with germ cell cyst breakdown (Pepling, 2006, 2012; Pepling and Spradling, 2001). Oogonium, before differentiating into oocytes, go through several mitotic divisions in which cytokinesis is not completed. Thus, resulting in a syncitium of cells, the cysts, that will initiate meiosis. At the end of the meiotic prophase, these cysts need to be broken so single oocytes can be surrounded by stromal ovarian cells to form follicles. The significance of proper cyst breakdown is manifested in mutants that fail to individualize oocytes, which are infertile due to the formation of follicles containing multiple oocytes (Xu and Gridley, 2013). Our model predicted that at least in part, oocyte loss would facilitate cyst breakdown. So, we expected that in mutants that fail to activate the DDR, cyst breakdown and follicle formation would be altered. Our data suggest that in our colony less than 45% of the oocytes of 15.5 dpc mice are forming cysts. However, this seems unlikely since cyst breakdown should not start until 17.5 dpc (Pepling and Spradling, 2001). Nonetheless, some reports show the existence of cyst fragmentation during fetal development that may result in the individualization of oocytes before 17.5 dpc (Lei and Spradling, 2013). However, this event seems to be rare during fetal development, since less than 7% of the oocytes were individualized at 17.5 dpc and this number only reach 20% by birth (Lei and Spradling, 2013). Thus, it is more likely that our data reflects that the approach we used for this analysis is unable to detect all oocytes in cysts. This is not unlikely especially because of the two-dimensional analysis we have performed, we were only able to determine if one oocyte was in a cyst if its sister cells were on the same plane of the section. Thus, all oocytes in cyst in which their sister cells are located in different planes may have been scored as individual oocytes. This will ultimately cause an overrepresentation of this category in our dataset.

Taking this caveat into consideration, to address if the DDR is required to breakdown cysts and if this had an effect on follicle formation, firstly we will focus our analysis in the follicle formation rate. The analysis of the *Chk2*^*-/-*^ ovaries shows that CHK2 does not participate in cyst breakdown since the number and proportion of follicles formed in *Chk2* mutants are indistinguishable to those found in control samples. Interestingly, in vitro inhibition of CHK1 function in *Chk2* mutant ovaries reduced the proportion of follicles formed in vitro (Table S5). Furthermore, these samples contained more oocytes in cysts, suggesting that the DDR was required for cyst breakdown. In our opinion, these data point out the importance of a functional DDR in the proper timing of follicle formation. These data suggest that DDR-dependent oocyte elimination may participate in cyst breakdown and follicle formation. In this sense, in *Spo11*^*-/-*^ ovaries, *Chk2* mutation specifically rescues the number of oocytes in cysts. Supporting the idea that the DDR impacts cyst breakdown, and consequently follicle formation.

Altogether, our data highlights the importance of the DDR in regulating the oocyte reserve in the females. First, during fetal development regulating the number of fetal oocytes using a CHK2-dependent response and postnatally activating a CHK1-dependent response. The oocyte loss caused by these two mechanisms promotes cyst breakdown, facilitating follicle formation and thus, setting the reserve of oocytes that mammalian females will use during their entire reproductive lifespan.

## Acknowledgements

We thank Attila Toth and Ewelina Bolcun-Filas for discussions and sharing unpublished information. We thank members of the SCAC for technical advice in setting up the ovarian culture technique. This work was supported by the Ministerio de Ciencia e Innovación (BFU2013-43965-P and BFU2016-80370-P, IR). AMM was supported by a PIF fellowship from Universitat Autònoma de Barcelona (B16P0048).

## Authors contributions

A.M.M. and I.R. designed the experiments. A.M.M., M.T.G., M.F. and A.G. performed experiments. A.M.M., M.T.G., M.F., A.G., M.G.C. and I.R. analyzed the data. A.M.M. and I.R. wrote the the paper with imput from all co-authors.

## STAR Methods

### Animals and genotyping

*Chk2* and *Spo11* mutant mice were generated previously (Baudat et al., 2000; Takai et al., 2002). These alleles were maintained in a C57BL6-129Sv mixed background. All experiments were performed using at least two mutant animals and compared with control littermates or with animals of closely related parents. The term wild type (WT) in the text and figures refers to both homozygous and heterozygous mice. All animals were sacrificed using CO_2_ or decapitation methods following the protocol CEEAAH 1091 (DAAM6395) approved by the Ethics Committee of the Universitat Autònoma de Barcelona and the Catalan Government.

Mouse genotyping was performed by PCR analysis from the DNA extracted from the tails as previously performed (Marcet-Ortega et al., 2017; Pacheco et al., 2015, 2018).

### Obtaining of the ovarian samples

Ovarian samples were obtained from females at different ages from 15.5 dpc to 22.5 dpc. The presence of a vaginal plug was used to determine the age of the unborn fetuses. Females were caged overnight with males and if the vaginal plug was found next morning the female was isolated and defined as 0.5 dpc. To obtain 15.5 and 17.5 dpc ovaries, pregnant females were sacrificed and the fetuses were removed, rinsed in PBS and ovaries were harvested under a stereo microscope (Nikon SMZ-1). Since in our mouse colony deliveries always occurred at 19.5 dpc, neonatal ovaries from 1 to 4 dpp correspond to 19.5 to 22.5 dpc.

### Histology, immunolabeling and oocyte quantification

Harvested perinatal ovaries (from 15.5 to 22.5 dpc) were immediately fixed overnight in 4% paraformaldehyde in PBS. Samples were then dehydrated, cleared and embedded in paraffin using standard procedures. The whole ovary was sectioned at 7 µm thickness and half ovary (every other section) was processed for immunostaining as follows: the sections were deparaffinized and antigen was unmasked by treating the slides for 20 minutes in Sodium Citrate buffer (10 mM Sodium Citrate, 0.05% Tween 20 in Milli-Q water, pH 6.0) at 95-100ºC. Standard immunofluorescence was then performed using rabbit anti-DDX4 (Abcam) at 1:100 dilution (Roig et al., 2004). Slides were counterstained with DAPI.

Oocytes were manually counted and classified under the fluorescence microscope (Zeiss Axipohot) with the 63x objective. Classification of the analyzed oocytes was done as follows: oocytes in cyst (sharing the cytoplasm, Fig. 3A), single oocytes (individual oocytes showing clear cytoplasmic limits, Fig. 3B) and oocytes in follicles (oocytes surrounded with one layer of flat granulosa cells, Fig. 3C). Only DDX4-positive oocytes with a visible nucleus stained with DAPI were considered for the analysis.

### Surface oocyte spreads and immunolabeling

Isolated ovaries from 17.5 and 19.5 dpc fetuses were placed in a 24-well plate with M2 medium (Sigma-Aldrich) and digested in collagenase dissolved in M2 medium for 20 minutes at 37ºC. Then the ovaries were transferred to hypotonic buffer (30 mM Tris-HCl pH=8.2, 50 mM Sucrose, 17 mM Sodium Citrate, 5 mM EDTA, 0.5 mM DTT [Roche Diagnostics], 1x Protease Inhibitory Cocktail [Roche Diagnostics] in Milli-Q water) and incubated for 30 minutes at room temperature. The oocytes were released from the ovary by pipetting up and down in 100 mM sucrose and transferred onto slides with fixative (1% PFA, 5 mM Sodium Borate, 0.15% Triton X-100, 3 mM DTT, 1x Protease Inhibitory Cocktail in Milli-Q water, pH= 9.2) in a humid chamber. After 2 hours of fixation, slides were air dried, washed with 0.4% Photoflo (Kodak), air dried again and stored at -80ºC until use.

Immunostaining was performed using standard methods (Roig et al., 2004) with the following primary antibodies: rabbit anti-SYCP3 (Abcam) at 1:200 dilution and mouse anti-γH2AX (Millipore) 1:200 dilution.

### Neonatal ovarian organ culture

Neonatal mouse ovaries were harvested and cultured, as previously reported (Morgan et al., 2015). Briefly, ovaries were placed in prewarmed dissection media (Leibowitz L15 [Sigma-Aldrich] with 3mg/mL of BSA) and under the stereo microscope, they were trimmed with forceps, eliminating the bursal sac, the oviduct and the uterus. Ovaries were transferred to UV-sterilized polycarbonate membrane (Whatman) sitting on a 24-well plate containing prewarmed culture media (α-Minimum Essential Medium (Thermofisher) with 3mg/mL of BSA). The samples were treated with 1 μM or 5 μM of the CHK1 inhibitor LY2603618 (Selleckchem), dissolved in DMSO. Control samples were treated with equivalent volumes of DMSO. All ovaries were cultured in an incubator at 37°C supplied with 5% CO_2_ for 5 days. After this time, ovaries were processed as previously mentioned to obtain ovarian sections.

### AZT treatment

To inhibit LINE-1 activity, 5mg/kg/day and 15 mg/kg/day AZT (Sigma Aldrich) were administered to wild-type and *Chk2*^*-/-*^ pregnant mice from 11.5 dpc until 17.5 dpc by gavage. At 17.5 dpc pregnant mice were sacrificed and fetal ovaries were obtained and processed as mentioned above to obtain ovarian sections.

### Image processing and data analysis

Microscopy analysis was performed with a Zeiss Axiophot microscope. Images were captured with a Point Gray Research, Inc. camera with the ACO XY Software (A. COLOMA Open microscopy).

All the images were processed with Adobe Photoshop CC to overlay the different fluorescent channels and match the intensity observed in the microscope. The software Image Composite Editor (https://www.microsoft.com/en-us/research/product/computational-photography-applications/image-composite-editor/) was used to stitch the composite panoramic images of the sections.

### Statistical analysis

Data analysis and statistical inference were performed using the GraphPad Prism 5 software (https://www.graphpad.com/scientific-software/prism/).

## Supplementary Information

**Figure S1. The organotypic culture allows folliculogenesis initiation *in vitro*.**

(A) Number of oocytes classified in the three different types (oocytes in cyst, single oocytes and oocytes in follicle) for control ovaries cultured for different number of days. After 5 days of culture, the ovaries present the same percentage of follicles as the 21.5 dpc ovaries (3 dpp).

(B) Histological section of a control ovary after 5 days of culture, immunostained against DDX4 and counterstained with DAPI. Scale bar represents 40 μm.

**Table S1.** Number of γH2AX patches per oocyte at pachynema and diplonema from WT and *Chk2*^*-/-*^ ovaries.

| Stage | WT | *Chk2*<sup>-/-</sup> |
| --- | --- | --- |
| Early pachynema | 52.9 ± 1.8* (N=50) | 59.3 ± 1.6* (N=50) |
| Late pachynema | 25.3 ± 1.1* (N=75) | 33.3 ± 1.1* (N=75) |
| Early diplonema | 14.9 ± 0.8* (N=36) | 17.5 ± 0.9* (N=36) |
| Late diplonema | 4.7 ± 0.6 (N=39) | 4.4 ± 0.5 (N=39) |

The numbers express the average ± SEM.
N indicates the number of oocytes counted.
\*represents the statistical difference between the two genotypes (T-test).

**Table S2.**
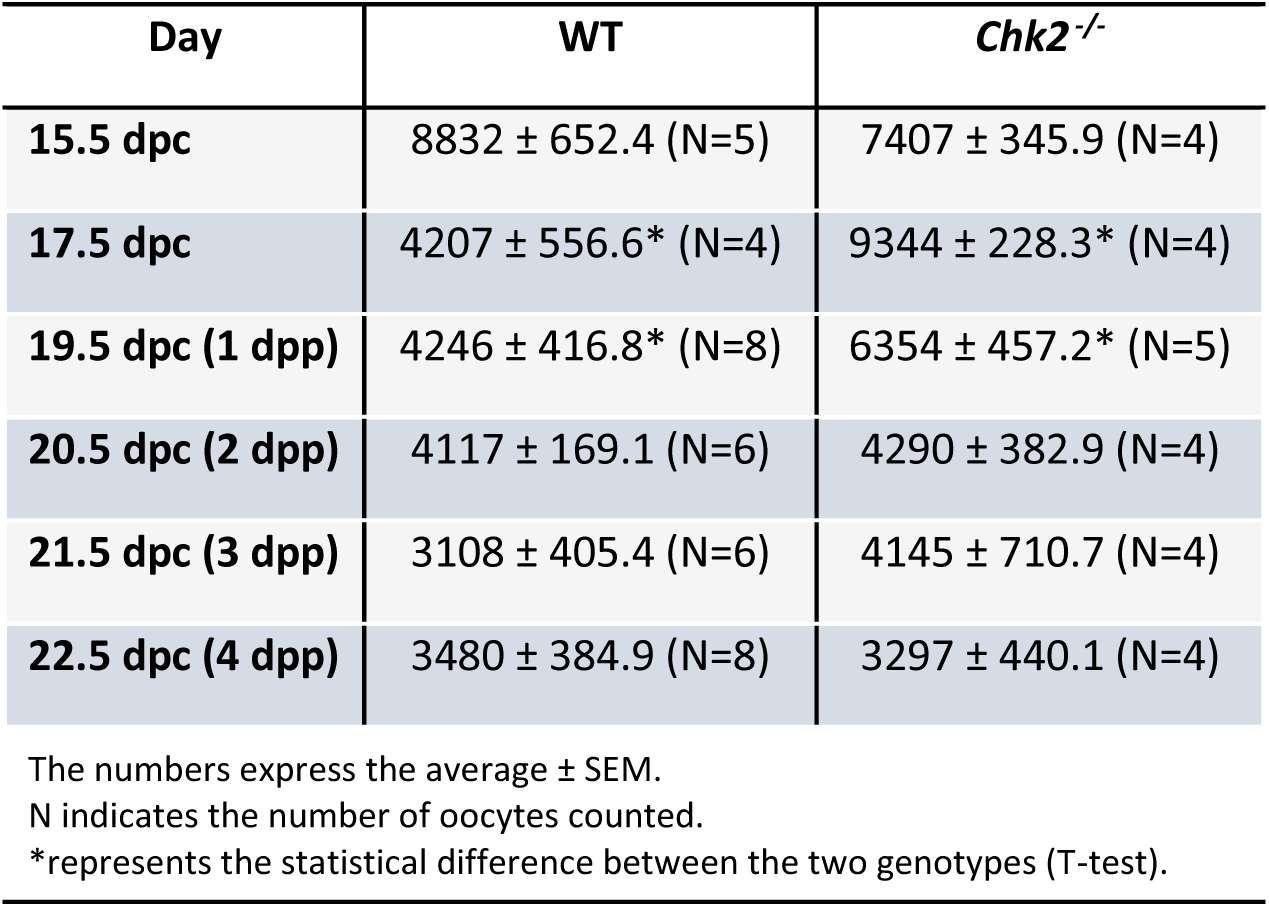
Oocyte number on the different perinatal days from WT and *Chk2*^-/-^ ovaries.

**Table S3.** Number and percentage of the different classes of oocytes found on the WT and *Chk2*^-/-^ ovaries analyzed.

| Day | Genotype |  | Oocytes in cyst | Single oocytes | Oocytes in follicles |
| --- | --- | --- | --- | --- | --- |
| 15.5 dpc | WT (N=5) | # | 3844 ± 335.3 | 4987 ± 398.8 | 0 |
|  |  | % | 43.6 ± 2 | 56.4 ± 2 | 0 |
|  | *Chk2*<sup>-/-</sup> (N=4) | # | 3057 ± 210.5 | 4351 ± 251.7 | 0 |
|  |  | % | 41.2 ± 2 | 58.8 ± 2 | 0 |
| 17.5 dpc | WT (N=4) | # | 1620 ± 236.2* | 2539 ± 310.9* | 48.6 ± 18 |
|  |  | % | 38.2 ± 1.4 | 60.7 ± 1.4 | 1.1 ± 0.3* |
|  | *Chk2*<sup>-/-</sup> (N=4) | # | 3591 ± 242.9* | 5735 ± 105.8* | 19.25 ± 7.5 |
|  |  | % | 38.3 ± 1.8 | 61.5 ± 1.8 | 0.2 ± 0.1* |
| 19.5 dpc<br>(1 dpp) | WT (N=8) | # | 1456 ± 203.6 | 2466 ± 323.9* | 324.3 ± 46.8 |
|  |  | % | 34.3 ± 4.1 | 57.9 ± 3.7 | 7.8 ± 1.1 |
|  | *Chk2*<sup>-/-</sup> (N=5) | # | 1875 ± 372.7 | 3929 ± 283.2* | 550.8 ± 119.8 |
|  |  | % | 28.9 ± 4.6 | 62.7 ± 5.5 | 8.3 ± 1.3 |
| 20.5 dpc<br>(2 dpp) | WT (N=6) | # | 677.7 ± 86.2 | 2675 ± 207.1 | 763.8 ± 117.5 |
|  |  | % | 16.5 ± 2 | 64.9 ± 3.9 | 18.6 ± 3 |
|  | *Chk2*<sup>-/-</sup> (N=4) | # | 593.8 ± 90.9 | 3005 ± 202.3 | 691.4 ± 269.7 |
|  |  | % | 14.6 ± 3.3 | 70.5 ± 2.7 | 14.9 ± 5 |
| 21.5 dpc<br>(3 dpp) | WT (N=6) | # | 167.7 ± 32.07 | 1212 ± 149.4 | 1728 ± 271.2 |
|  |  | % | 5.6 ± 0.9 | 40 ± 3.1 | 54.5 ± 3.7 |
|  | *Chk2*<sup>-/-</sup> (N=4) | # | 304.1 ± 118.4 | 1308 ± 153 | 2533 ± 458.1 |
|  |  | % | 6.4 ± 1.8 | 32.9 ± 3.5 | 60.7 ± 2.6 |
| 22.5 dpc<br>(4 dpp) | WT (N=8) | # | 163.9 ± 47.9 | 1028 ± 158.2 | 2288 ± 205.2 |
|  |  | % | 4.2 ± 0.8 | 29 ± 1.8 | 66.8 ± 2.4 |
|  | *Chk2*<sup>-/-</sup> (N=4) | # | 74 ± 14.07 | 1040 ± 84.3 | 2183 ± 381.7 |
|  |  | % | 2.3 ± 0.4 | 32.6 ± 3.3 | 65.1 ± 3.6 |

The numbers express the average ± SEM.
N indicates the number of oocytes counted.
\*represents the statistical difference between the two genotypes (T-test).

**Table S4.**
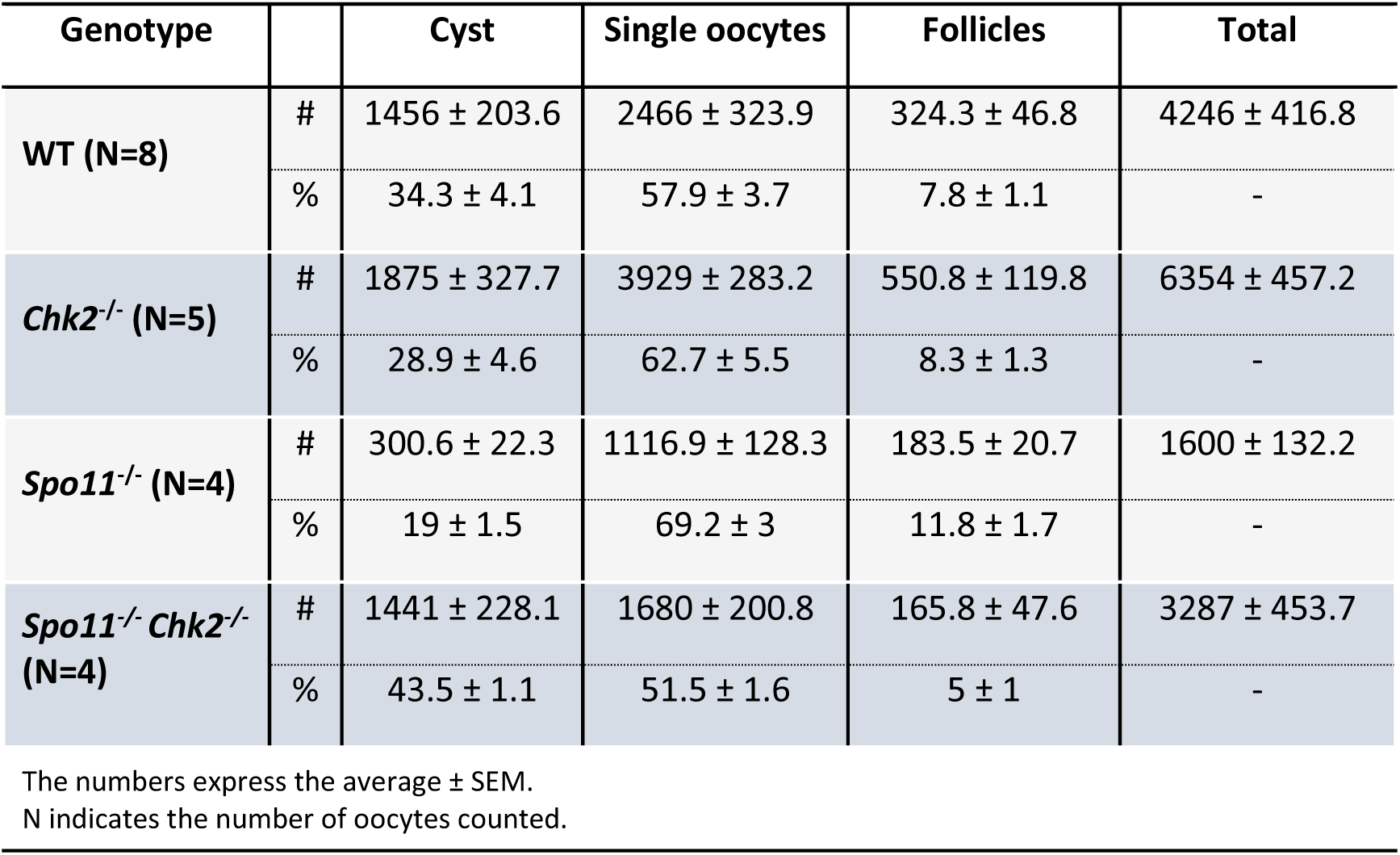
Number and percentage of the different classes of oocytes found at 19.5 dpc ovaries from the different indicated genotypes.

| Genotype |  | Cyst | Single oocytes | Follicles | Total |
| --- | --- | --- | --- | --- | --- |
| **WT (N=8)** | # | 1456 ± 203.6 | 2466 ± 323.9 | 324.3 ± 46.8 | 4246 ± 416.8 |
|  | % | 34.3 ± 4.1 | 57.9 ± 3.7 | 7.8 ± 1.1 | - |
| ***Chk2*<sup>-/-</sup> (N=5)** | # | 1875 ± 327.7 | 3929 ± 283.2 | 550.8 ± 119.8 | 6354 ± 457.2 |
|  | % | 28.9 ± 4.6 | 62.7 ± 5.5 | 8.3 ± 1.3 | - |
| ***Spo11*<sup>-/-</sup> (N=4)** | # | 300.6 ± 22.3 | 1116.9 ± 128.3 | 183.5 ± 20.7 | 1600 ± 132.2 |
|  | % | 19 ± 1.5 | 69.2 ± 3 | 11.8 ± 1.7 | - |
| ***Spo11*<sup>-/-</sup> *Chk2*<sup>-/-</sup> (N=4)** | # | 1441 ± 228.1 | 1680 ± 200.8 | 165.8 ± 47.6 | 3287 ± 453.7 |
|  | % | 43.5 ± 1.1 | 51.5 ± 1.6 | 5 ± 1 | - |

The numbers express the average ± SEM. N indicates the number of oocytes counted.

**Table S5.** Number and percentage of the different classes of oocytes found at 19.5 dpc *Chk2*^*-/-*^ cultured ovaries at the indicated conditions.

| Genotype |  | Cyst | Single oocytes | Follicles | Total |
| --- | --- | --- | --- | --- | --- |
| **DMSO (N=9)** | # | 94.6 ± 25.7 | 559.8 ± 46.7 | 824.8 ± 67.6 | 1479 ± 124.4 |
|  | % | 5.9 ± 1.2 | 38.1 ± 1.5 | 56 ± 1.9 | - |
| **Chk1i 1 μM (N=3)** | # | 116.8 ± 48.6 | 467.3 ± 187.5 | 848.5 ± 168.7 | 1433 ± 401.3 |
|  | % | 7.4 ± 2.2 | 30.3 ± 4.1 | 62.3 ± 5.8 | - |
| **Chk1i 5 μM (N=6)** | # | 320.6 ± 73.4 | 1033 ± 134.2 | 693.8 ± 95.5 | 2047 ± 260.5 |
|  | % | 14.9 ± 1.9 | 50.6 ± 2.6 | 34.4 ± 3.7 | - |

The numbers express the average ± SEM. N indicates the number of oocytes counted.

**Table S6.** Number and percentage of the different classes of oocytes found at 18 dpc *Chk2*^*-/-*^ AZT treated ovaries at the indicated conditions.

| Day | Genotype |  | Cyst | Single oocytes | Follicles | Total |
| --- | --- | --- | --- | --- | --- | --- |
| Water | WT (N=7) | # | 496 ± 34.2* | 3268 ± 303.6* | 54.4 ± 11 | 3818 ± 299* |
|  |  | % | 13.5 ± 1.4 | 85 ± 1.7 | 1.6 ± 0.4 | - |
|  | *Chk2*<sup>-/-</sup> (N=6) | # | 901.2 ± 28.8* | 6207 ± 423.4* | 68.7 ± 12.1 | 7177 ± 454.5* |
|  |  | % | 12.7 ± 0.6 | 86.4 ± 0.5 | 0.9 ± 0.1 | - |
| AZT 5 mg/kg | WT (N=6) | # | 495.7 ± 64.3 | 2704 ± 256.8 | 14.7 ± 6.2 | 3215 ± 249.8 |
|  |  | % | 16 ± 2.6 | 83.6 ± 2.5 | 0.4 ± 0.2 | - |
|  | *Chk2*<sup>-/-</sup> (N=4) | # | 867.8 ± 290.6 | 5641 ± 1654 | 58.5 ± 25.2 | 6567 ± 1964 |
|  |  | % | 12.8 ± 1.3 | 86.5 ± 1.3 | 0.8 ± 0.2 | - |
| AZT 15 mg/kg | WT (N=3) | # | 724.7 ± 121.3 | 4685 ± 197.3 | 82 ± 10* | 5492 ± 206.3 |
|  |  | % | 13.2 ± 2.1 | 85.3 ± 2.1 | 1.5 ± 0.2* | - |
|  | *Chk2*<sup>-/-</sup> (N=6) | # | 844.3 ± 50 | 6537 ± 563.2 | 48.7 ± 6.4* | 7430 ± 599.2 |
|  |  | % | 11.5 ± 0.6 | 87.8 ± 0.6 | 0.7 ± 0.1* | - |

The numbers express the average ± SEM.
N indicates the number of oocytes counted.
\*represents the statistical difference between the two genotypes (T-test).

