## Supplementary figures and images for "The DNA damage response regulates the oocyte pool in mammals"

### Fig S1

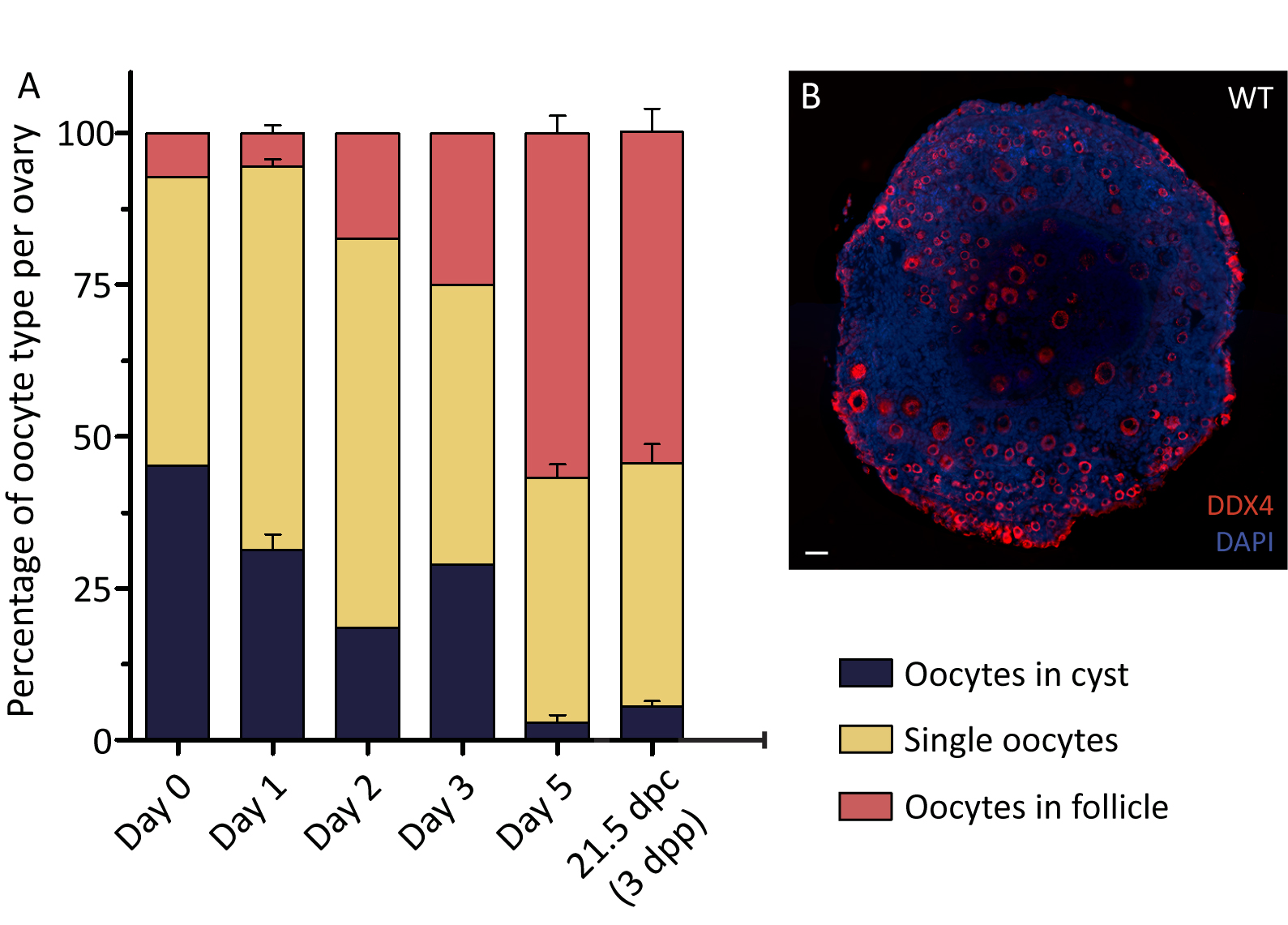
